# The antigen presenting molecule MR1 binds riboflavin catabolites

**DOI:** 10.1101/2025.07.20.665728

**Authors:** Mohamed R. Abdelaal, Jieru Deng, Mitchell P. McInerney, Emi Ito, Anthony W. Purcell, Sho Yamasaki, Jose Villadangos, Hamish E. G. McWilliam, Nicholas A. Gherardin, Jamie Rossjohn, Wael Awad

## Abstract

Major histocompatibility-complex (MHC) class I-related (MR1) protein presents vitamin B based antigens to Mucosal-Associated Invariant T (MAIT) cells. While microbial riboflavin precursors are well documented MR1 ligands, it is unclear whether host-generated riboflavin catabolites influence MR1-mediated immunity. Here, we report that riboflavin catabolites, including 10-formylmethylflavin (FMF), lumichrome, lumiflavin and alloxazine bind to MR1 with moderate affinity, while riboflavin itself binds weakly. In contrast to the microbial riboflavin antigens which increase MR1 cell surface expression, the riboflavin catabolites moderately reduced cell surface levels of MR1 by stabilizing and retaining MR1 in the endoplasmic reticulum (ER). The riboflavin catabolites appeared to bind to the intracellular MR1 and inhibit MR1 exit from the ER. These riboflavin catabolites also weakly competed with Vit B based Ags for MR1 binding, thereby inhibiting MAIT cell activation. The crystal structures of MR1 complexed with riboflavin, FMF, lumichrome and lumiflavin, show binding of these three-ringed ligands in the A’-pocket of MR1. The crystal structure of MR1-lumichrome revealed that lumichrome formed a covalent “flavin bond” with MR1-Lys43 differing from the typical Schiff-base bond of MR1-Lys43-Ag complexes. Collectively, we identified three ring isoalloxazines that can bind MR1 and downregulate cell surface expression levels, suggesting a potential role in dampening MAIT cell immunity.

## Introduction

The major histocompatibility complex (MHC) I-related molecule, MR1, captures and presents small organic molecules to an innate-like T cell population called mucosal-associated invariant T (MAIT) cells (Awad et al., 2023; Corbett et al., 2014; Kjer-Nielsen et al., 2012). MAIT cells play crucial roles in both protective and aberrant immunity, as well as mucosal homeostasis (Corbett et al., 2020; Provine and Klenerman, 2020). MR1 is ubiquitously expressed in all cells but its cell surface expression is dependent on ligand availability. The MR1 ligand-binding A′-pocket has adequate malleability to capture a broad range of chemical structures including vitamin B antigens (VitBAg) exemplified by vitamin B9 derivatives (6-formylpterin (6-FP) and acetyl-6-FP (Ac-6-FP)) (Eckle et al., 2014; Kjer-Nielsen et al., 2012) and vitamin B6 derivatives (pyridoxal and pyridoxal-5’-phosphate) (McInerney et al., 2024), as well as non-VitBAg compounds like environmental ligands (e.g., components of cigarette smoke (CS)) (Awad et al., 2025), drugs and drug-like molecules and other small molecule compounds (Keller et al., 2017; Salio et al., 2020; Wang et al., 2022a). More recently, host-derived nucleobase adducts, and sulfated bile acids have been described (Chancellor et al., 2025; Ito et al., 2024; Vacchini et al., 2024). MR1 also presents microbial derivatives of vitamin B2 (riboflavin; RF) precursors that are formed during infection with a wide range of RF-producing microbes (Corbett et al., 2014). Here, the microbial intermediate 5-amino-6-D-ribitylaminouracil (5-A-RU) non-enzymatically reacts with small glycolysis metabolites (e.g., methylglyoxal) to generate short-lived intermediates e.g. 5-(2-oxopropylideneamino)-6-D-ribitylaminouracil (5-OP-RU) that are captured by MR1 before conversion to lumazines e.g. RL-6-Me-7-OH, and are potent antigens (Ags) for MAIT cells. These RF-based Ags function as a “*microbial metabolite signature*” that activate MAIT cells, where 5-OP-RU represents the most potent MAIT cell agonist identified to date (Awad et al., 2020a; Awad et al., 2020b). However, no host-derived riboflavin-related catabolites have been described.

Typically, the binding of MR1 to ligands within the endoplasmic reticulum (ER) of antigen-presenting cells (APCs) triggers ER-resident MR1 to translocate to the cell surface (McWilliam et al., 2016; McWilliam et al., 2020). MR1-ligand complexes remain at the cell surface for several hours before re-internalized back to the cell interior and/or endosomes (McWilliam et al., 2016), with the majority of MR1 molecules subsequently being degraded, and a minor fraction recycling from endosomes back to the cell surface, potentially loaded with a different ligand. The release of MR1 from the ER to the cell surface involves a conformational change driven by a molecular switch involving a lysine residue (Lys-43) at the base of the ligand-binding groove. Here, the potent MR1-binding ligands form a covalent bond, termed a “Schiff base” with Lys-43, neutralizing its positive charge and stabilizing MR1 (McWilliam and Villadangos, 2024). Indeed, mutation of MR1 Lys-43 to Ala (termed MR1^K43A^) prevents Schiff base formation with these ligands, dramatically reducing Ag-presentation (McWilliam et al., 2016; Reantragoon et al., 2013). Notably, some synthetic ligands, such as DB28 and NV18.1 compounds that bind MR1 but do not form Schiff base bonds, can retain MR1 within the ER, leading to a downregulation at the cell surface (Salio et al., 2020), but whether naturally occurring ligands can suppress MR1 upregulation remains unclear.

A number of reports have provided evidence that RF itself cannot activate MAIT cells, but rather blocks their response to bacterial antigen presentation, reminiscent of other non-stimulatory MR1-binding ligands such as 6-FP (Harriff et al., 2018; Kjer-Nielsen et al., 2012; Shibata et al., 2022). The mechanisms of RF-based MAIT cell antagonism, however, are unknown. When RF is catabolized (enzymatically or by photodegradation) in humans, distinct products are formed including: 10-Formylmethylflavin (FMF), lumiflavin, lumichrome (7,8-dimethylalloxazine) and/or alloxazine, (Bitsch and Bitsch, 2016), all of which can be detected in human blood (Barupal and Fiehn, 2019; Hardwick et al., 2004). Given that these RF catabolites are structurally similar to the microbial MR1 binding Ags, we reasoned that MR1 may bind these host RF catabolites. Through cellular, biochemical, and structural approaches, we show that RF catabolites bind MR1 and impact its cell surface levels, thereby showing how host derived three ring-compounds can modulate the MR1-MAIT axis.

## Results

### Riboflavin and RF catabolites bind MR1 molecules

To investigate whether RF and its catabolites **(Fig. 1A)** are potential ligands for MR1, we quantified the relative binding affinities of these compounds for MR1 using our established fluorescence polarization (FP)-based cell-free assay (Wang et al., 2022a). The IC_50_ concentrations were interpolated from the non-linear regression of the dose-response curves for MR1-ligand interaction (**Fig. 1B**). Consistent with previous reports (Wang et al., 2022a), 5-OP-RU and Ac-6-FP, which form a Schiff base interaction with MR1, show strong binding to MR1 with IC_50_ values: 5.3 and 29.9 nM, respectively. The FP assay showed that RF is a weak binder to MR1 with an IC_50_ value of ∼182 µM, consistent with the previous finding that RF can stabilize MR1 on the cell surface of antigen presenting cells (Shibata et al., 2022). However, when RF was exposed to UV light (30 min) or daylight (6-8 h) [referred as photodegraded RF hereafter], the IC_50_ reduced to ∼45 µM, suggesting the formation of higher affinity MR1 ligands in solution during RF photodegradation, reminiscent of folic acid photodegradation products having MR1-binding capacity (Kjer-Nielsen et al., 2012). We then measured the IC_50_ for individual RF catabolites (**Fig. 1B**), which were moderate binders with IC_50_ values in the range 12 – 54 µM. Mass spectrometry analysis showed that MR1-photodegraded RF contained a mixture of compounds including lumichrome (MH+ as m/z 243.087) and carboxymethylflavin (MH+ as m/z 301.093) **(Fig. 1C-G)**.

**Figure 1:**
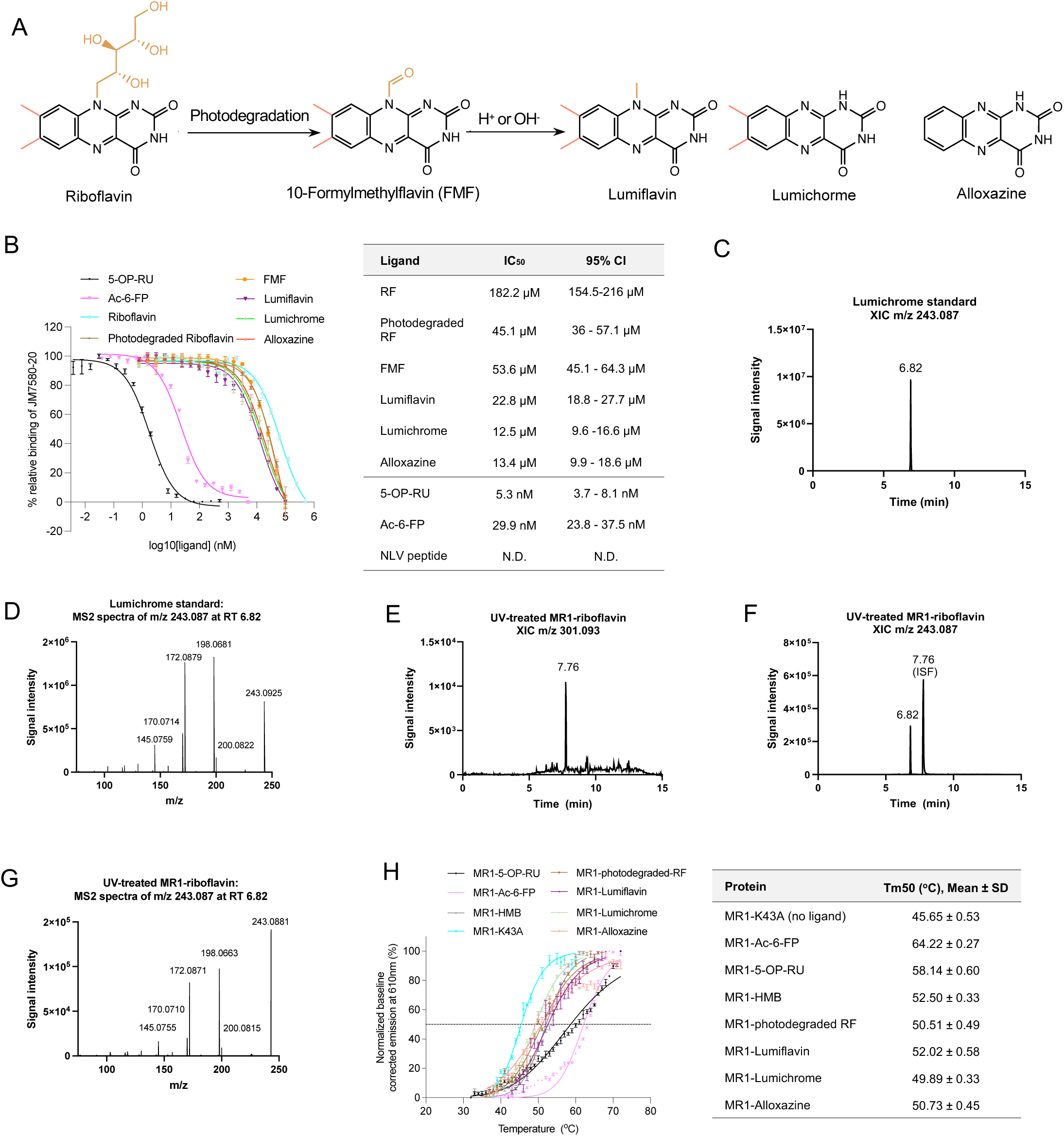
Riboflavin and riboflavin catabolites bind MR1. **(a)** Mechanism of metabolism and photodegradation of riboflavin. **(b)** Titration curves of the shown ligands binding to MR1 were obtained from fluorescence polarization (FP)-based assay (left). Each data point represents normalized percentage binding from two independent experiments performed in triplicate. Mean values are plotted with SEM represented in error bars. Curve fit for ligands are displayed in the table (right). **(c)** The extracted ion chromatograms (XICs) for m/z 243.087 in a lumichrome solvent standard and **(d)** MS2 fragmentation of main peak at RT 6.82 are depicted. **(e)** UV-treated MR1 riboflavin XIC for m/z 301.0927. **(f)** XIC for m/z 243.087 showing lumichrome present at RT 6.82, with an additional peak at RT 7.76. **(g)** MS2 fragmentation of main peak at RT 6.82, showing aligned fingerprint comparison with lumichrome solvent standard. **(h)** Thermostability of soluble WT MR1 refolded with the indicated ligands was measured by fluorescence-based thermal shift assay. The graph shows baseline-corrected, normalized emission at 610 nm plotted against temperature (□). Each point represents the mean of three technical replicates and error bars represent SD. The half maximum melting point (Tm50) is indicated by the dotted line at 50%. The table on the right shows the mean Tm50 from three independent experiments, each measured in at least a technical triplicate. RF; riboflavin.

We then assessed the capacity of RF and its catabolites to be loaded within recombinant MR1. The human MR1-β2-microglobulin (β2m) complex can only co-purify when the MR1 Ag-binding pocket is occupied with ligand (Kjer-Nielsen et al., 2012). Photodegraded RF, as well as purified FMF, lumiflavin, lumichrome and alloxazine facilitated the correct folding of the MR1-β2m co-complex in solution while RF did not sponsor refolding of MR1 (**Fig. S1A-B**). Next, we examined how these catabolites could stabilize MR1 using an *in vitro* MR1-Ag complexes through thermostability assay (Eckle et al., 2014) (**Fig. 1H**). Here, the half-maximum melting temperatures (Tm50) of the stable complexes MR1-5-OP-RU and MR1-Ac-6-FP were 58.65 ± 0.42 and 62.12 ± 0.37 °C; respectively, consistent with previous values (Eckle et al., 2014). MR1^K43A^-β2m, which refolds freely of Ag, cannot form a Schiff base and is considered to be unstable, had a Tm50 of 45.97 ± 0.5 °C. The RF catabolites had a moderate effect on stability of MR1 where MR1-Photodegraded RF, MR1-lumiflavin, MR1-lumichrome and MR1-alloxazine complexes had Tm50 values ranging from 49.9-52.0□°C. Thus, the binding of these ligands stabilizes the MR1 protein but to a lower extent in comparison to MR1-5-OP-RU and MR1-Ac-6-FP (**Fig. 1H**).

### RF-based catabolites down-regulate MR1 cell surface expression

Next, we investigated whether these RF catabolites can impact MR1 cell surface expression in both MR1-transduced B cell lymphoblastoid (C1R.MR1) and monocytic (THP-1.MR1) cell lines (**Fig. 2**). Here, the cells were pulsed with 1-200 µM of RF or the RF catabolites, and the cell surface levels of MR1 were quantified by flow cytometry (Awad et al., 2020a; Keller et al., 2017). Cell viability was unaffected, even at highest concentrations of the catabolites, as depicted in **Fig. S2A**. In C1R.MR1 cells, RF -up to 200 µM-did not alter MR1 expression from the baseline after 3 h (**Fig. 2A**) but modestly upregulated MR1 after 16 h (**Fig. 2B-C**), whereas none of the RF catabolites could up-regulate MR1 cell surface expression at either timepoint. Instead, lumichrome and alloxazine reduced MR1 cell surface after 3 h (**Fig. 2A**) while all RF catabolites downregulated MR1 after 16 h in a concentration-dependent manner (**Fig. 2B-C**). No impact on the MR1 surface level was observed when the parental C1R cells were incubated with these RF catabolites, probably due to the low basal surface MR1 level whose further reduction becomes undetectable by the anti-MR1 antibody (**Fig. S2B**). The MR1 downregulation seen in C1R.MR1 cells was specific to MR1 as the level of HLA-A*02:01 was not affected after pulsing C1R-A2 with the ligands (**Fig. S2C**). A similar trend was observed after pulsing THP-1-MR1 with RF catabolites for 3 and 16 h **(Fig. 2D-2F)**, yet lumiflavin downregulated MR1 after 3 h (**Fig. 2D**), in comparison to C1R.MR1 cells. This MR1 downregulation varied between cell types, with a higher extent observed in THP-1.MR1 than C1R.MR1, and was seen after staining with both two conformational anti-MR1 8F2F9 and 26.5 antibodies (clones 8F2F9 and 26.5; **Fig. 2G)**. Overall, our data suggest that the direct interactions of RF catabolites with MR1 in the cells lead to the reduction of cell surface expression where lumichrome is the most potent down-regulator followed by alloxazine, lumiflavin and then FMF.

**Figure 2.**
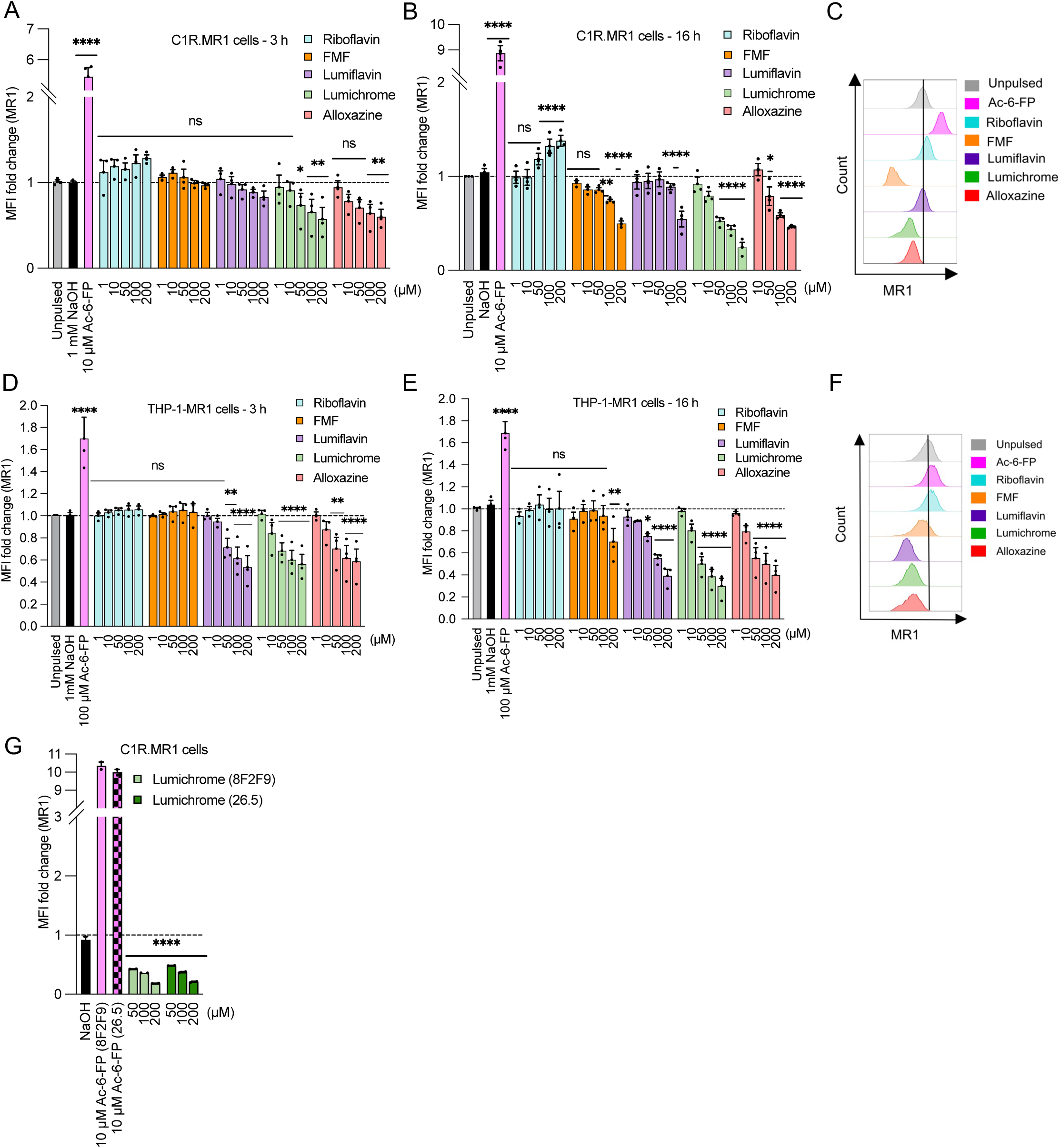
Riboflavin catabolites modulate MR1 expression on the surface of antigen presenting cells. **(a-b)** Bar graphs depict the expression of surface MR1*01 on C1R.MR1 cells incubated for 3 h **(a)** or 16 h **(b)** with titrated quantities of ligands followed by staining with 8F2F9 anti-MR1 antibody. **(c)** Overlay histograms showing the expression of surface MR1*01 on C1R.MR1 cells after 16 h treatment with the indicated concentrations of RF catabolites. **(d-e)** MR1 was then quantified after adding the indicted concentrations of the riboflavin catabolites on THP-1.MR1 cells incubated for 3 h **(d)** or 16 h **(e)**. **(f)** Overlay histograms showing the expression of surface MR1*01 on THP-1.MR1 cells after 16 h treatment with the indicated concentrations of RF catabolites. **(g)** The bars indicate the expression of surface MR1 on C1R.MR1 cells after treatment with lumichrome overnight and then staining with 26.5 vs. 8F2F9 MAb for comparison. Data represent the geometric mean fluorescence intensity (gMFI) fold change from three independent experiments performed in duplicates, with standard error (SEM) represented by the error bars. One-way ANOVA statistical analysis was performed for all samples with multiple comparisons performed using NaOH as a control (ns: not significant, * *p* <0.05, \*\**p* < 0.01, \*\*\**p* < 0.001, \*\*\**p* < 0.0001). MR1, MHC-I related protein 1.

### MR1-binding RF catabolites keep MR1 in the ER in an immature state

The downregulation of MR1 on the cell surface by the RF catabolites could be due to the depletion of the intracellular ER-resident MR1 pool, or by retention of MR1 in the ER, as shown for the DB28 ligand (Salio et al., 2020). To investigate this, C1R.MR1 cells were treated for 16 h with 100 µM of RF or RF catabolites, conditions which induce MR1 downregulation (**Fig. 3A**). Cells were lysed, and treated with endoglycosidase (Endo) H, to distinguish molecules that remain within the ER from those that have trafficked to the cell surface (McWilliam et al., 2016), then MR1 levels were assessed by immunoblotting (**Fig. 3A**). In the absence of exogenous ligand, the majority of MR1 remains predominantly endo H-sensitive, as demonstrated by the dominant band of lower molecular weight on SDS-PAGE. Addition of Ac-6-FP, which triggers MR1 egression from the ER, results in an increase in MR1 of a higher molecular weight, in line with Endo H-resistance (**Fig. 3A**). Neither RF nor its catabolites induced any Endo H-resistant MR1; however, RF catabolites consistently increased the level of Endo H-sensitive MR1 molecules. This indicates that these RF catabolites do not deplete ER-resident MR1 but rather retain and induce its accumulation.

**Figure 3.**
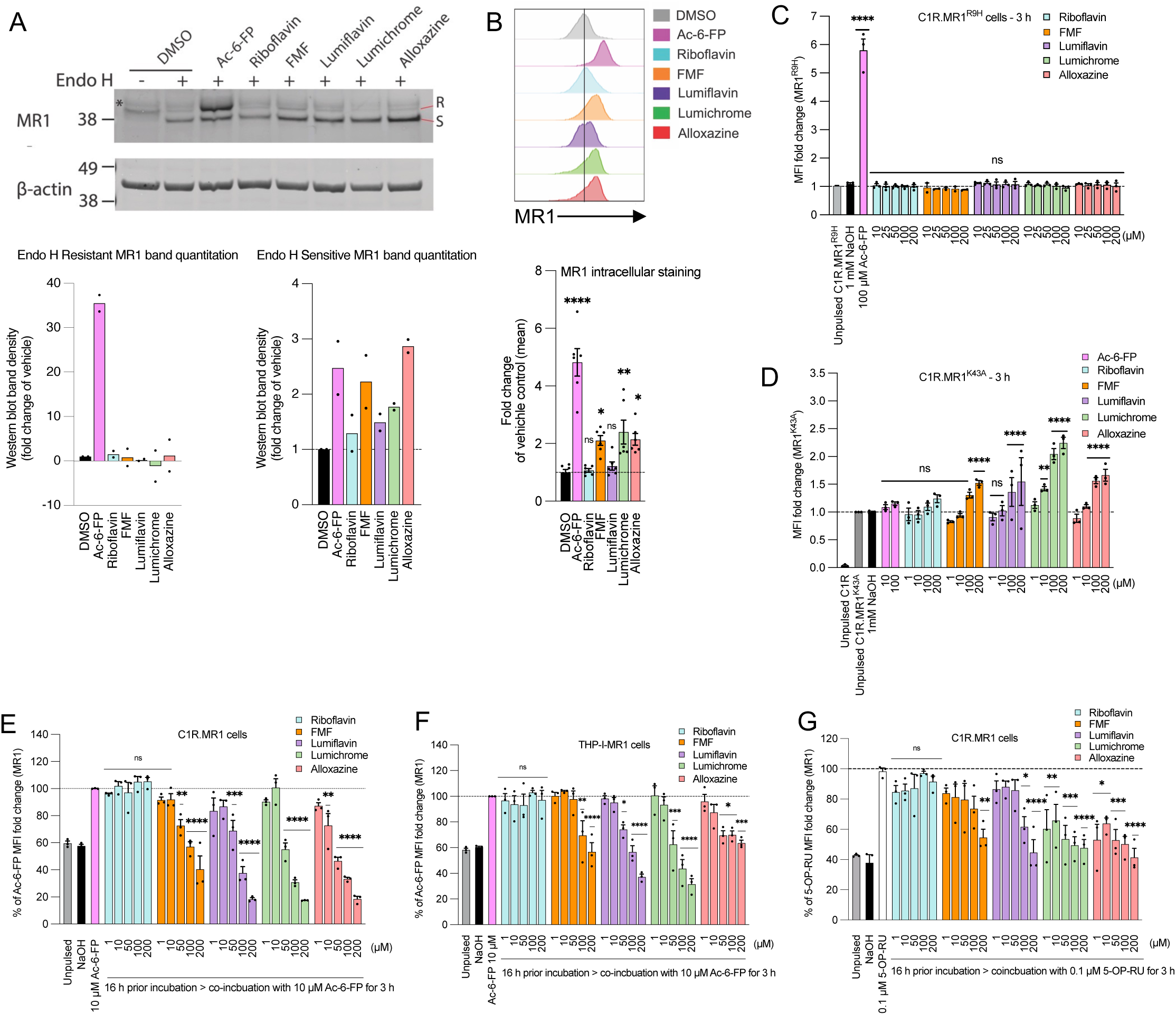
Riboflavin catabolites induce retention of MR1*01 in the ER, but not MR1^K43A^. **(a)** Analysis of Endo-H-treated (+) or untreated (-) MR1 by western blotting with anti-MR1 (8G3) after culturing C1R.MR1 cells with DMSO (vehicle control), Ac-6-FP (10μM), riboflavin (100μM), FMF (100μM), lumiflavin (100μM), lumichrome (100μM) or alloxazine (100μM) for 16 h, at 37□. S, Endo H–susceptible MR1; R, Endo H–resistant MR1. The Endo H–susceptible MR1 and Endo H–resistant MR1 fractions were quantified in at least two independent experiments. **(b)** Intracellular total MR1 level was also measured in C1R.MR1 cells by flow cytometry after treating the cells with the indicated ligands (100 µM) for 16h followed by permeabilization and staining with anti-MR1-PE (8F2.F9). Shown are the overlay histograms **(top)** and a bar chart depicting the geometric mean fluorescence intensity (gMFI) fold change of intracellular MR1 level **(bottom)** from three independent experiments performed in duplicates, with standard error (SEM) represented by the error bars. C1R cells expressing **(c)** MR1^R9H^ mutant or **(d)** MR1^K43A^ mutant were incubated for the indicated periods with titrated quantities of ligand followed by flow cytometry. **(e-g)** The competition between RF catabolites and Ac-6-FP in **(e)** C1R.MR1 cells and **(f)** THP-I.MR1 cells as well as **(g)** 5-OP-RU in C1R.MR1 cells was quantified after pulsing with the indicated concentrations of RF catabolites for 16 h before the addition of Ac-6-FP/5-OP-RU for further 3 h. Shown in (**e-g)** is the average percentage reduction in Ac-6-FP/5-OP-RU-induced MR1 upregulation in three independent experiments performed in duplicates with standard error (SEM) represented by error bars. One-way ANOVA statistical analysis was performed for all samples with multiple comparisons performed using NaOH as a control (ns: not significant, * *p* <0.05, \*\**p* < 0.01, \*\*\**p* < 0.001, \*\*\**p* < 0.0001). MR1, MHC-I related protein 1.

Previously, it was shown that within the ER, there are two forms of MR1 – ‘folded’ and ‘open’ – and these can be distinguished with a conformationally-sensitive antibody 8F2.F9, which specifically recognises the ‘folded’ form (McWilliam et al., 2016; McWilliam et al., 2020). The ‘open’ MR1 are not bound to β2m, whereas the folded MR1 is associated with β2m, are more stable and show less degradation in the absence of ligands (McWilliam et al., 2016; McWilliam et al., 2020). To determine if the RF catabolites increase the level of either forms of ER-resident MR1 molecules, C1R.MR1 cells were treated with or without RF and its catabolites as above, and the levels of folded MR1 detected by intracellular staining with 8F2.F9 and detected by flow cytometry **(Fig. 3B)**. As expected, Ac-6-FP significantly increased total folded MR1, which includes both intracellular and cell-surface complexes. The RF catabolites FMF, lumichrome and alloxazine, but not with RF or lumiflavin, also increased the total level of folded MR1 (**Fig. 3B**).

This suggests that FMF, lumichrome and alloxazine induce the ‘folded’ conformation of intracellular MR1 molecules. This is consistent with the hypothesis that these metabolites bind to ‘open’ ER-resident MR1 molecules and induce their folding to the more stable ‘closed’ conformation. This stabilises these complexes without inducing their release from this compartment, leading to their intracellular accumulation. This provides a mechanistic explanation for the observed downregulation of MR1 from the cell surface in the presence of these metabolites.

### Requirements for RF catabolites-mediated down-modulation of MR1 cell surface levels

We then tested the effect of the RF catabolites on the expression of two mutant versions of MR1 termed MR1^R9H^ (Howson et al., 2020) and MR1^K43A^ (Reantragoon et al., 2013). First, we tested the MR1^R9H^ mutant (Howson et al., 2020) which represents a rare, natural human polymorph. It lacks the ability to bind and present RF metabolites such as 5-OP-RU, yet retains the ability to bind other ligands such as folate metabolite Ac-6-FP, offering an opportunity to further understand the molecular constraints on MR1 binding to RF derivatives. As expected, the level of MR1^R9H^ on the surface of C1R cell transfected with MR1^R9H^ (C1R.MR1^R9H^) significantly increased after Ac-6-FP treatment (**Fig. 3C**). When C1R. MR1^R9H^ were incubated with RF and RF catabolites, MR1^R9H^ surface level did not deviate from the background level (**Fig. 3C**). In addition, lumichrome did not refold with hMR1^R9H^ in solution. Thus, similar to microbial RF-derivate ligands, RF catabolites also do not bind MR1^R9H^. Next, we tested sought to understand how the previously characterised Lys43A mutation affects ligand binding. This mutation stabilizes MR1 and helps refolding in the absence of exogenous ligands (Reantragoon et al., 2013) and thus the level of trafficked MR1^K43A^ is significantly increased on C1R.MR1^K43A^ cell surface (**Fig. 3D**). Surprisingly, lumichrome, alloxazine and lumiflavin increased MR1^K43A^ at the cell surface after 3 h (**Fig. 3D**) and this MR1 upregulation continues with lumichrome as well as RF after 16 h (**Fig. S2D**). While this is in line with the ability of these ligands to bind MR1, the reason for MR1^K43A^ cell surface accumulation is less clear. The most likely explanation is that these ligands bind ER-resident MR1^K43A^, in line with our data using wild-type MR1; however, because the K43A mutation results in spontaneous MR1 surface egress, the presence of these stabilising ligands promotes enhanced trafficking and/or retention at the cell surface. We cannot rule out however that the ligands may also bind directly to surface MR1^K43A^ and slow down the recycling from cell surface.

### MR1-binding RF catabolites compete with VitBAgs for MR1 binding

We next tested whether FMF, lumiflavin, lumichrome and alloxazine impact the MR1 cell surface expression induced by Ac-6-FP or 5-OP-RU. The RF catabolites were incubated with C1R.MR1 and THP-1.MR1 cells overnight at 1-200 µM concentration before the addition of either 10 µM Ac-6-FP or 5-OP-RU, then co-incubated for another 3 h. RF itself did not modulate Ac-6-FP or 5-OP-RU responses **(Fig. 3E-G)**. In comparison, the four RF catabolites reduced the Ac-6-FP-induced cell surface upregulation of MR1 in a concentration-dependent manner, more profoundly in C1R.MR1 (**Fig. 3E**) than THP-1.MR1 cells (**Fig. 3F**). Interestingly, the highest concentration (200 µM) of lumiflavin, lumichrome and alloxazine could reduce more than 80% of Ac-6-FP response. The competition with 5-OP-RU was less potent with high concentrations of the RF catabolites in C1R.MR1 cells **(Fig. 3G)**, in line with the stronger affinity of 5-OP-RU for MR1, compared to Ac-6-FP, as measured by the FP assay (**Fig. 1B**). This cellular data, along with the moderate affinity for MR1 revealed by the FP assay, suggests that FMF, lumiflavin, lumichrome and alloxazine are able to bind MR1 molecule within cells and hindering the capacity of other potent ligands to stimulate MR1 trafficking to the cell surface.

### MR1-binding RF catabolites impact MAIT cell effector functions *in vitro*

Next, we explored whether RF catabolites could activate SKW-3 cells expressing two different MAIT TCRs. Distinct responses of T cells expressing TCRs with varying TRBV chain usage reinforces the role of V_β_ in modulating MAIT responses to various ligands (Chancellor et al., 2023; Gherardin et al., 2016; Seneviratna et al., 2022).The surface expression of CD69 was measured as a marker for activation of cells expressing the AF-7 (TRAV1-2/TRBV6-1) or MBV28 (TRAV1-2/TRBV28) TCRs after co-culture with C1R.MR1 cells and the ligands. As expected, 5-OP-RU induced strong activation of both cell lines while 6-FP moderately reduced basal CD69 levels (**Fig. 4A-B and S4B-S4C**) (Eckle et al., 2014). While minor reductions in CD69 were seen at the highest dose of lumichrome for both TCRs, and a minor increase in CD69 for AF-7 in response to riboflavin and lumiflavin, these fluctuations were subtle relative to the potency of 5-OP-RU (**Fig. 4A-B and S4B-S4C**).

**Figure 4.**
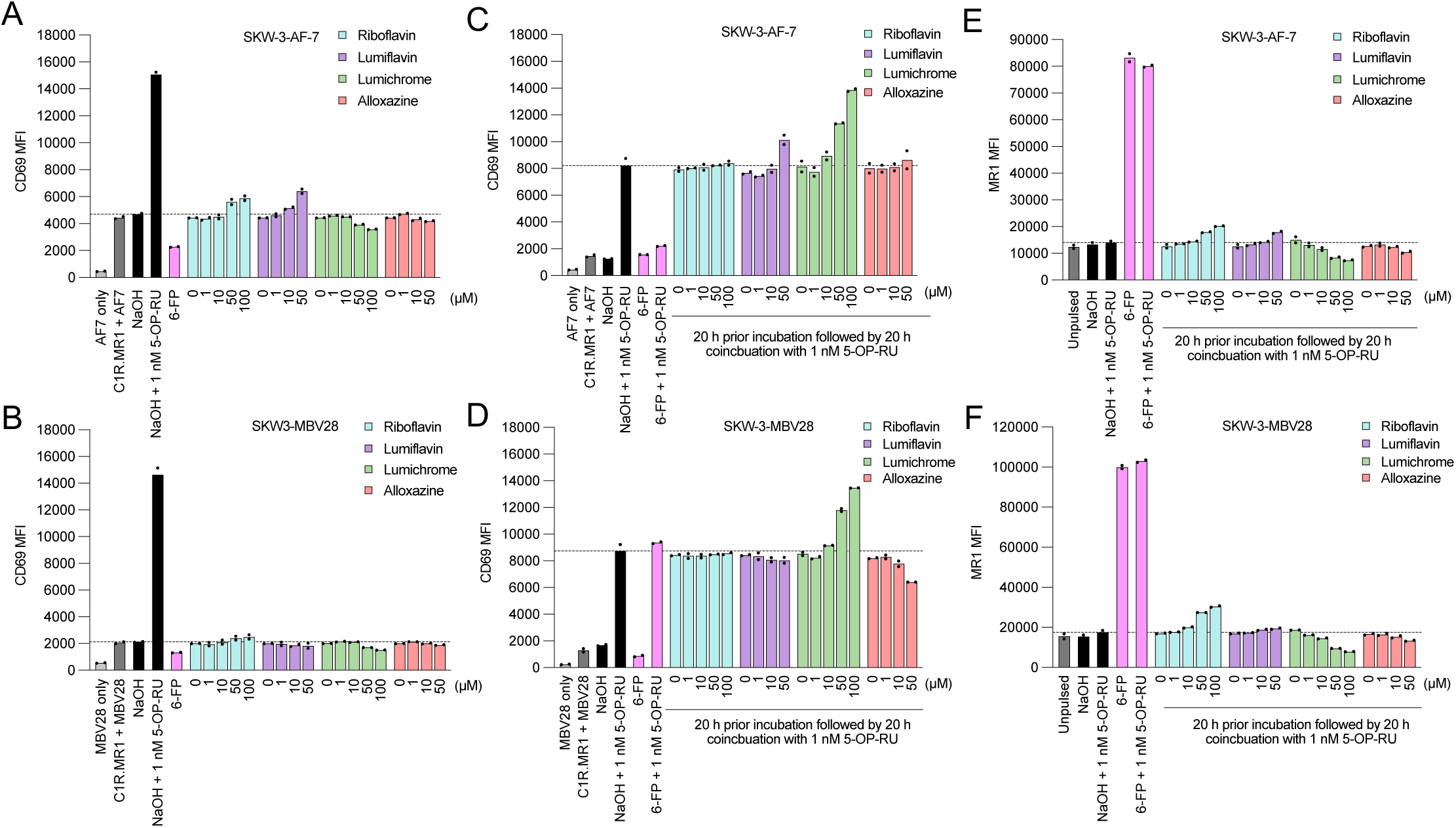
Riboflavin-based catabolites impact on MAIT cell activation. Bar graphs showing CD69 expression on two separate SKW-3-MAIT cell lines; clones AF-7 (TRAV1-2^+^TRBV6-1^+^) and MBV28 (TRAV1-2^+^TRBV28^+^) cultures in the absence **(a-b)** or presence of 5-OP-RU **(c-d)** after preincubation with titrated quantities of indicated ligands followed by harvesting, staining and flow cytometric analysis. The MR1 expression on C1R.MR1 cell surface within the (c-d) 5-OP-RU co-incubation experiments is shown in **(e-f)**. Shown here is a representative of at least 3 independent experiments. Individual datapoints are technical replicates from a representative experiment.

When the T cells were incubated with RF catabolites followed by co-incubation with 5-OP-RU, differential responses in both MR1T cell clones were observed (**Fig. 4C-D and S4D-E**). RF could not compete with 5-OP-RU for the activation of either SKW-3-AF-7 nor SKW-3-MBV28 cells. High concentration of lumiflavin slightly exaggerated the 5-OP-RU-induced activation of AF-7 (**Fig. 4C and S4D**), but not MBV28 (**Fig. 4D and S4E**). Alloxazine could interfere with the 5-OP-RU-induced activation of MBV28 in a dose-dependent manner, but not the AF-7 cells. In comparison, lumichrome enhanced the 5-OP-RU-induced CD69 expression on the cell surface of both cell lines (**Fig. 4C-D and S4D-E**). MR1 levels were also monitored in these co-cultures and, in agreement with **Fig. 2F**, we observed a dose-dependent reduction in MR1 expression after coincubation with lumichrome at the indicated concentrations (**Fig. 4E-F and S4F-G**). This might be due to the lumichrome-induced accumulation of MR1 in the ER, so the cells have more MR1 molecules to capture 5-OP-RU. Altogether, these data suggest that RF-derived catabolites bind MR1, and their relative effects on activation, either alone or in combination with 5-OP-RU, are varied and clone dependent.

### MR1-binding RF catabolites inhibit MAIT cell effector functions ex vivo

The data using C1R.MR1 and SKW-3 cells above gave unexpected results, whereby RF-catabolites variably modulated MAIT activation in a clone-dependent manner. This data is possibly confounded by the artificially high levels of MR1 both in the ER and on the surface, which may enhance opportunities for ligand exchange or other uncharacterised pathways of ligand presentation. Therefore, we next employed an assay system that uses primary APCs and MAIT cells which more closely mimic physiological antigen-presentation conditions. Here, we co-cultured healthy human PBMC with RF or it’s catabolites and assessed the effects on MAIT cell activation. First, MAIT proliferation was measured after CTV-labeled PBMCs were co-cultured with various RF ligands. CD3+MR1-5-OP-RU tet+ MAIT cells significantly proliferated upon stimulation with 5-OP-RU. Lumichrome significantly reduced MAIT cell proliferation in comparison to the other RF catabolites which did not significantly change the baseline level (**Fig. 5A and Fig. S3D**). MAIT activation was then evaluated by quantifying CD69 expression on the surface of gated MAIT cells after 16-h incubation with RF and its catabolites. While 5-OP-RU induced robust CD69 upregulation, none of the RF catabolites induced CD69 upregulation beyond that of known MAIT antagonist Ac-6-FP (**Fig. 5B**). Furthermore, unlike the SKW-3.MAIT-based system, upon co-incubation with 5-OP-RU, RF and all RF catabolites showed potent concentration-dependent competition with 5-OP-RU leading to consequent MAIT inhibition (**Fig. 5C**). Lumichrome was the most potent competitor to 5-OP-RU. This data showed that MR1-binding RF catabolites, especially lumichrome, can bind to MR1 and act as antagonists for MAIT cells.

**Figure 5.**
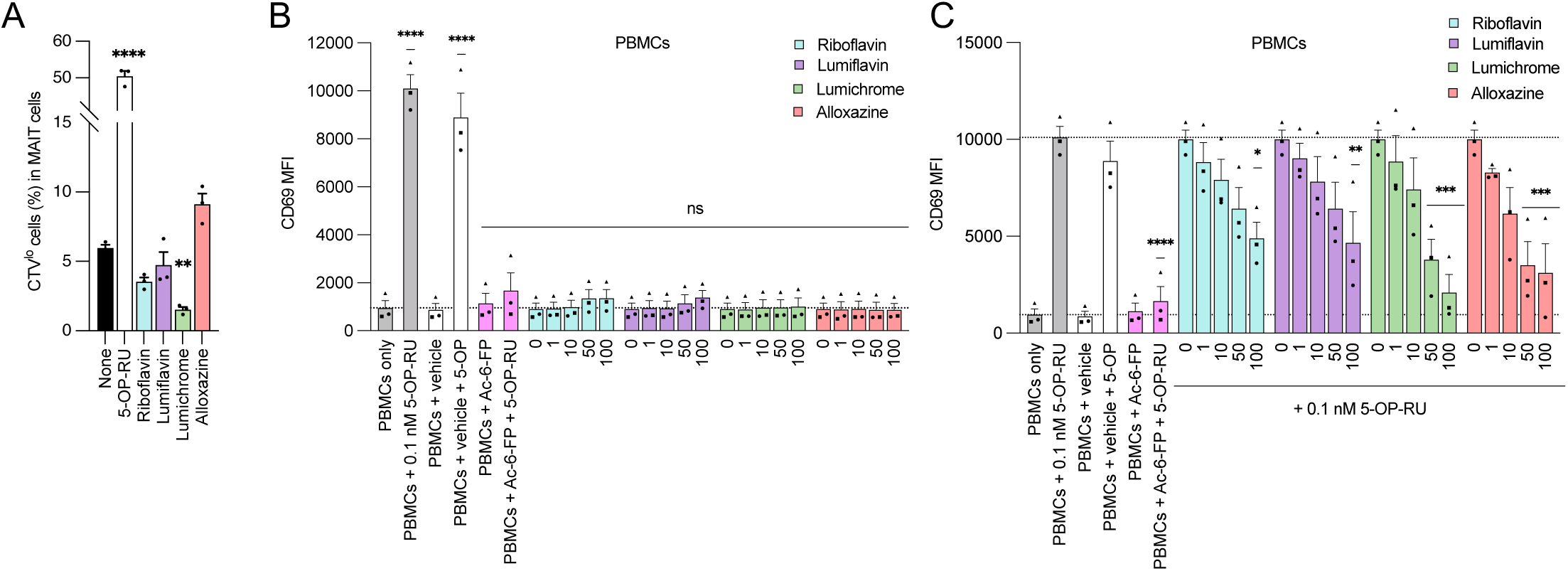
Riboflavin-based catabolites impact on *ex-vivo* MAIT cell activity in PBMCs. **(a)** Human PBMCs were labeled with Cell Trace Violet (CTV) and stimulated with vehicle control (None), 5-OP-RU (10 µM), riboflavin (100 µM), lumiflavin (100 µM), lumichrome (100 µM) or alloxazine (100 µM) on day 7. Proportion of MR1−5-OP-RU tet^+^CTV^lo^ MAIT cells in CD3^+^5-OP-RUtet^+^-gated cells was shown. Data shown in **a** is the average of at least three independent experiments performed in duplicates with standard error (SEM) represented by error bars. **(b-c)** Depicted is the average CD69 expression in MAIT cells gated from three donors measured after titration with the indicated RF ligands for 16 h, in the absence **(b)** or presence **(c)** of 5-O-RU, followed by flow cytometry. Data in **b-c** represent average median fluorescence intensity (MFI) from three donors performed in duplicates, with standard error (SEM) represented by the error bars. One-way ANOVA statistical analysis was performed for all samples with multiple comparisons performed using NaOH as a control for **a and b** and 0.1 nM 5-OP-RU as a control for **c** (ns: not significant, * *p* <0.05, \*\**p* < 0.01, \*\*\**p* < 0.001, \*\*\**p* < 0.0001)

### Crystal structures of MR1-RF-derived ligands

To understand the structural basis for the binding of RF and RF catabolites to MR1, we determined the high-resolution crystal structures of MR1 in complex with RF, photodegraded-RF, FMF, lumiflavin and lumichrome (**Fig. 6** and **Table S1**). Crystallization of MR1-binary structures is challenging, so we used the A-F7 MAIT TCR (Tilloy et al., 1999) to aid crystallization, as reported previously (McInerney et al., 2024; Salio et al., 2020). In the MR1-photodegraded RF, the electron density within the MR1 A′-pocket is large and suggests the presence of more than one ligand (**Fig. S1C**), in line with the mass spectrometry data, so we did not refine further. The AF-7 TCR acquired typical docking mode previously seen in MR1-ligand-AF-7 ternary structures (**Fig. 6A**). The electron density for the remaining ligands within the MR1 A′-pocket were unambiguous (**Fig. 6B-E**), thereby permitting detailed structural analysis. In accommodating these three-ringed ligands (**Fig. 6F-I**), minimal conformational changes within the pocket residues were observed, and the side chains of the MR1 antigen-binding residues were mostly conserved. Notably, lumichrome was the only ligand that formed a covalent interaction with MR1-Lys43 (**Fig. 6B-E**).

**Figure 6.**
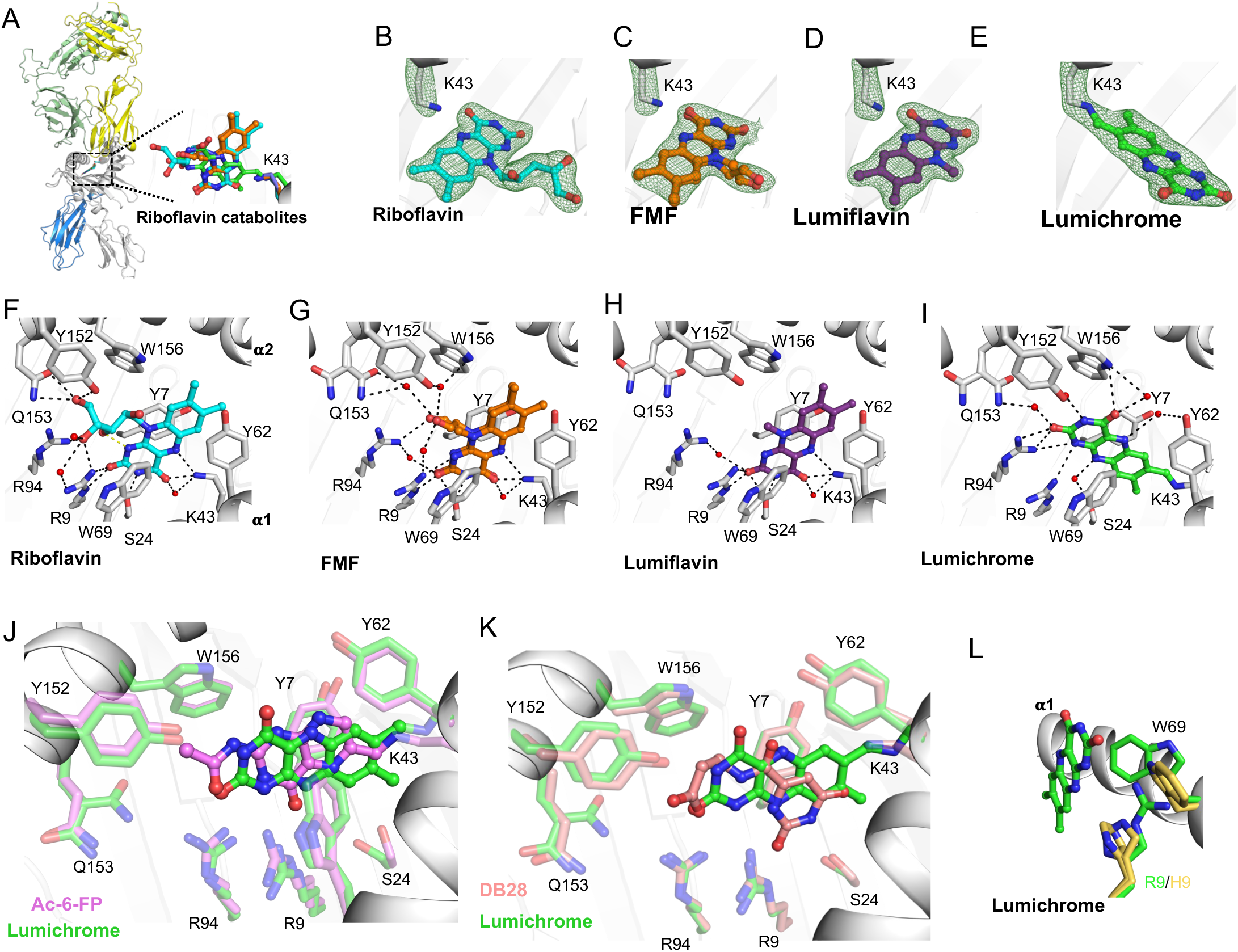
Overall docking and molecular interactions of riboflavin and RF catabolites within MR1 A’-pocket. **(a)** Superposition of the TCR-MR1-riboflavin catabolites crystal structures showing the riboflavin and its catabolites within the MR1-binding cleft of the MR1-riboflavin-AF-7 structure interacting with MR1-Lys43. **(b-e)** The electron density omit maps (green mesh) of **(b)** riboflavin, **(c)** FMF, **(d)** lumiflavin and **(e)** lumichrome contoured at 2σ. **(f-I)** The molecular contacts of **(f)** riboflavin, **(g)** FMF, **(h)** lumiflavin and **(i)** lumichrome with the residues of MR1-A′ pocket in the MR1-Ag structures. **(j-k)** Superposition of lumichrome (green) with **(j)** Ac-6-FP (pink; PDB: 4PJ5) and **(k)** DB28 (salmon; PDB: 6PVC) within MR1 ligand-binding cleft. **(l)** Superposition of Lumichrome within the MR1 ligand-binding pocket of MR1^R9H^-AF-7 structure (PDB; 6W9V) showing the MR1 residues from MR1-lumichrome-AF-7 structure in green and MR1^R9H^ -AF-7 residues in yellow. MR1 and β2-microglobulin (β2m) are coloured white and marine, respectively. Here, the cut-off for hydrogen bonds, salt bridges and VDW interactions were set at 3.5 Å, 4 Å and 4 Å, respectively. Ligands are coloured as follows: riboflavin, cyan; FMF, orange; lumiflavin, magenta; and lumichrome, green. The α and β chains of the AF-7 TCR are coloured yellow and pale green, respectively.

In the crystal structure of MR1-RF, RF did not form a Schiff base with MR1-Lys43 (**Fig. 6B**); yet the 4-carbonyl and 5-nitrogen in the flavin (isoalloxazine) ring formed H-bonds with Lys43 (**Fig. 6F**). The 2-carbonyl and 3-nitrogen groups of the RF ring formed H-bonds with MR1-Arg9 and Ser24, respectively. The ribityl moiety formed H-bonds with MR1-Arg9, Arg94, Tyr152 and Gln153, in addition to VDW interaction with MR1-Trp156 (**Fig. 6F**). The positioning of the ribityl tail is governed by a network of intramolecular and intermolecular polar contacts. The intramolecular interaction is formed between 3′-OH group and the isoalloxazine ring (**Fig. 6F**). In addition, Tyr152 and Gln153 of MR1 formed hydrogen bonds with the 5′-OH group.

In the crystal structures of MR1-FMF and MR1-lumiflavin, the flavin rings of both FMF and lumiflavin ligands had the same orientation as for RF and as such was involved in a similar network of interactions (**Fig. 6G-H**). The acetyl moiety in FMF adopted two conformations in the pocket. This acetyl moiety allowed form H-bond with MR1-Arg94 as well as water-mediated interactions with the α2 residues Tyr152, Gln153 and Trp156 (**Fig. 6G**). Here, the lack of ribityl tail in both FMF and lumiflavin disfavoured the formation of ‘direct’ hydrogen bonding with MR1-α2 residues (**Fig. 6G-H**). This shows clearly that RF catabolites are well-positioned within the MR1 A’-pocket, allowing the stabilization of MR1 in the ER and reduced trafficking to the cell surface.

### Lumichrome forms a “flavin bond” with MR1-Lys43

Lumichrome was sequestered within the MR1 aromatic cradle where its isoalloxazine ring sat in a perpendicular orientation to that of the others RF catabolites (**Fig. 6I**). This lumichrome ring orientation is similar to the potent MR1 up-regulators Ac-6-FP and 6-FP (Eckle et al., 2014) (**Fig. 6J**), ruling out that the downregulation observed with lumichrome is due to changes in lumichrome docking inside MR1. Here, the flavin ring of the lumichrome ligand was sandwiched between MR1-Tyr7, Tyr62, Trp69, and Trp156 while the 2-carbonyl moiety formed H-bonds with MR1-Arg94, water-mediated H-bonds with MR1-Gln153, as well as VDW interactions with MR1-Arg9 (**Fig. 6I**). Also, the 4-cabonyl moiety formed a direct H-bond with MR1-Trp156. The amino groups of the flavin ring formed direct H-bonds with MR1-Arg9 and Arg94 as well as water-mediated H-bonds with MR1-Ser24, Tyr62, and Try152. As lumichrome was unable to refold with MR1^R9H^ mutant, MR1-Arg9-lumichrome interaction is most likely essential for the ligand stabilization **(Fig. 6L)** and induced MR1 downregulation **(Fig. 3C)**. Surprisingly, a unique MR1-Lys43-flavin ring linkage “flavin bond” was observed between the 7-methyl group of the lumichrome ring and MR1-Lys43 (**Fig. 6I**). To compare, the MR1 ligand DB28 couldn’t form a covalent interaction with MR1-Lys43 (Salio et al., 2020), though the orientations of MR1 pocket residues is comparable to that of the lumichrome structure (**Fig. 6K**) explaining the similar downregulation trend. The mechanisms of covalent flavin incorporation (flavination) and the possible roles of covalent protein-flavin bonds has been previously reported in a significant number of flavoenzymes which use a covalently protein-bound flavin cofactor (Heuts et al., 2009; Wang et al., 2022b). A proposed mechanism for the covalent binding of lumichrome to MR1 is depicted in **Fig. S5.**

## Discussion

Riboflavin (vitamin B_2_) is essential for humans, as it is necessary for the synthesis of the flavin coenzymes, including flavin mononucleotide (FMN) and flavin adenine dinucleotide (FAD), which have key roles in oxidative metabolism. When RF is catabolized, distinct structurally-related products are formed according to the environment, whereby FMF and lumichrome are the major by-products under neutral or acidic pH conditions, while both lumiflavin and lumichrome are formed under basic pH (Bitsch and Bitsch, 2016). Indeed, human blood contains RF catabolites (Barupal and Fiehn, 2019; Hardwick et al., 2004). For example, the human plasma contains on average 24 and 11.5 nM RF and lumichrome, respectively (Hardwick et al., 2004). In addition, previous reports showed that high doses of RF catabolites prove efficacy against different types of cancer (Chantarawong et al., 2019; Machado et al., 2013; Yang et al., 2022) and osteolytic diseases (Liu et al., 2018); however, the immunological roles of the RF catabolites are yet unexplained. Here, we showed that RF, and a panel of RF catabolites (FMF, lumichrome and lumiflavin, and alloxazine) can bind and stabilize the antigen-presenting protein MR1. Further, when the cells were pulsed with RF catabolites, there was a notable reduction in MR1 surface expression with the majority of them. This can be attributed, at least in part, to the retention of the ligand-receptive MR1 molecules in the ER and thus preventing the trafficking to the cell surface. The induction of ‘folded’ MR1 and stabilisation is evidence that these compounds are binding within the ER. How they prevent MR1 egress to the cell surface is not clear.

All MR1 binding ligands characterized so far are accommodated within the A′ pocket. Many of these antigens stabilize MR1 by forming a Schiff base with Lys43 at the base of the pocket; however, this is not a strict requirement for MAIT cell activation. Here, we found that MR1 was anchored to lumichrome ligand by forming a “flavin bond” with MR1-Lys43. Although lumiflavin and FMF have a very similar chemical structure, they didn’t form this flavin bond and exhibited differential binding modes and contacts within the MR1 A′-pocket, with their isoalloxazine rings positioned closer to the MR1 α1 helix compared to the lumichrome. This unique interaction formed between lumichrome and MR1-Lys43 is an example for ‘covalent flavination’ of nonenzymatic immune-related proteins. Covalent flavination has nonetheless been reported more broadly, in particular in a large number of flavoenzymes which use a covalently protein-bound flavin cofactor (Heuts et al., 2009). This covalent tethering is a post-translational and self-catalytic process that was shown to tune the reactivity of the flavin group so that it can fulfil its catalytic role in the flavoproteins (Starbird et al., 2017; Wang et al., 2022b). Approximately 1% of all known proteins are flavoproteins (Piano et al., 2017) where about 10% of these flavoproteins contains a covalently bound flavin (Heuts et al., 2009). Being ubiquitous in all domains of life, they have extremely versatile catalysis activity in a broad range of essential biochemical processes, including natural product biosynthesis, photosynthesis, DNA damage repair, chromatin modification as well as immune-related activities (Piano et al., 2017; Teufel, 2017). Various immune-related proteins are covalently ‘flavinated’ where the attached flavin group(s) appear to be important for their function (McNeil et al., 2014; Moreno et al., 2022; Pedrolli and Mack, 2014; Zafred et al., 2013; Zhong et al., 2020). From a structural perspective, there are nine known types of flavin-protein linkages up to date, comprising His, Tyr, Cys, Asp, ser or Thr attached to either FAD, FMN or lumichrome (Wang et al., 2022b). In these linkages, the amino acid residue is attached to C6 or C8 of the flavin isoalloxazine ring or to the phosphate group. Here, we define a previously uncharacterized form of flavination in a non-enzymatic protein comprising the Lys-43 of MR1 covalently linked to C7 of the isoalloxazine ring. The biological significance of MR1 covalent flavination requires further investigation.

Downregulation of cell surface antigen-presenting proteins is a common strategy utilized by different microbes to evade MHC and CD1d-restricted T cell responses (Forsyth and Eisenlohr, 2016; Schuren et al., 2016; Van Kaer, 2007). Previously, the DB28 and its methyl analogue, NV18.1, were reported to downregulate the cell surface MR1 by retaining the MR1 molecules in the ER in an immature form and inhibit MAIT cell activation (Salio et al., 2020). However, these compounds are synthetic and do not occur in nature. Here, we report lumichrome, amongst other RF catabolites, as a host derived metabolite that specifically reduces the cell surface MR1 and competes with 5-OP-RU and Ac-6-FP for both ligand-induced MR1 upregulation as well as MAIT cell activation, thereby modulating MR1-MAIT signalling. This may represent a natural suppression mechanism to regulate cell-surface MR1 and to avoid inadvertent MAIT cell activation, thereby and providing control over the MR1-MAIT axis.

## Materials and methods

### Ligands

Riboflavin (Cat. No. R9504), lumichrome (Cat. No. 103217) and alloxazine (Cat. No. A28651) were purchased from Sigma-Aldrich. Lumiflavin was supplied by Cayman Chemical (Cat. No. 20645). Ac-6-FP (Cat. No. 11.418) and 6-FP (Cat. No. 11.415) were synthesized by Schircks Laboratories. JYM20 was synthesized as previously described (Wang et al., 2022a). All the ligands were dissolved in dilute NaOH to 10 mM and diluted when required. The solubilized ligands were always protected from lab light during storage and experiments.

To synthesise FMF, a solution of sodium periodate (2.2 g, 10 mmol) in water (23.5 mL) was added to a suspension of riboflavin (1.0 g, 2.7 mmol) in aqueous sulfuric acid (2 N, 26.5 mL) at 0 °C. After stirring at the same temperature for 0.5 h, the mixture was warmed to room temperature. After stirring at the same temperature for a further 16 h, the mixture was adjusted to pH 3.9 (as measured by a pH meter) with solid sodium carbonate. The precipitate was collected by filtration, and then washed successively with cold water, ethanol, and then diethyl ether. The precipitate was dried on high vacuum to give FMF as an orange solid (650 mg, 2.29 mmol, 85 %) as a mixture of the aldehyde and its corresponding hydrate. The NMR spectrum of FMF matched that reported in the literature. (Crielaard et al., 2022)

### Screening of the binding affinity of the ligands to MR1 using fluorescence polarization (FP) assay

Fluorescence polarization (FP)-based cell-free assay has been recently developed for quantitating the relative binding affinities of putative ligands for MR1, including both activators and inhibitors of MAIT cells (Wang et al., 2022a). This assay reflects the inverse correlation between the IC_50_ of a given ligands and the binding affinity of that ligand, the ability to form a Schiff base as well as the number of MR1/ligand non-covalent interactions. Various concentrations of RF and its catabolites were incubated in competition with 10 nM JYM20 (TAMRA-conjugated weak MR1 ligand) for binding 100 nM empty hMR1 protein in FP assay buffer (25 mM HEPES, pH 7.5, 150 mM NaCl, 5 mM EDTA) (Wang et al., 2022a). The fluorescence polarization of TAMRA was measured after 24 of incubation at 37°C using PHERAstar microplate reader (BMG LABTECH). The ligand-binding curves were simulated by non-linear regression with Prism software (GraphPad Software Inc.) using a sigmoidal dose-response curve. Here, the IC_50_ values reflect the binding affinity and are calculated as the ligand concentration required for 50% inhibition of JYM20 binding to MR1 molecules. The relative binding values (%) were then calculated as the percentage ration between IC_50_ value of the substituted JYM20 divided by the IC_50_ value of the non-substituted ligand at the 10 nM concentration.

### Quantification of cell surface MR1

The level of MR1 expressed on the cell surface was measured after the exposure to the investigated ligands as previously described (Awad et al., 2020a; Wang et al., 2022a). C1R antigen presenting B lymphocytes overexpressing MR1*01 (C1R.MR1), MR1^K43A^ mutant (C1R.MR1^K43A^), MR1^R9H^ mutant (C1R.MR1^R9H^) or HLA-A*02:01 (C1R-A2), in addition to the monocytic THP-1-MR1 cells, were used in this study. 2 x 10^5^ cells were incubated with RF catabolites for 3 or 16 h at 37° C and 5% CO2 in 200 μl folate-free RPMI 1640 medium from Gibco (Cat. No. 11875-093) fortified with 10% fetal bovine serum (FBS), sodium pyruvate (1 mmol/L), Hepes buffer (15 mmol/L), pH 7.2 to 7.5, 2% penicillin (100 U/ml), streptomycin (100 mg/ ml), glutamax (2 mmol/L), nonessential amino acids (0.1 mmol/L) (all from ThermoFisher, Life Technologies), and 2-mercaptoethanol (50 mmol/L, Sigma) (RF-10). After incubation, the cells were first stained with Zombie Aqua Fixable Viability Kit from BioLegend (Cat. No. 423102) and then incubated with biotinylated 8F2F9 αMR1 antibody for 30 minutes on ice. The unbound antibody was then washed off with 2% fetal bovine serum/PBS. The cells were finally incubated with PE-conjugated streptavidin (BioLegend, 30 μg/ml) and then washed with 2% fetal bovine serum/PBS. The PE fluorescence intensity reflects the surface MR1 level. The data was acquired with an LSRFortessa X-20 (BD Biosciences) and Diva software (BD Biosciences) and analyzed using FlowJo software (BD Biosciences). The gating strategy is shown in **Fig. S3A**.

### MAIT cell activation assay

C1R.MR1 cells (1 x 10^5^) were initially stained with CTV for 15 minutes and then incubated with RF catabolites for 20 h. SKW-3 cells (1 x 10^5^) overexpressing AF-7 or MBV28 MAIT TCRs were then tested for activation by co-incubation at a 1:1 ratio with the CIR.MR1 cells for further 20 h in 200□µL complete medium with various RF catabolites, in the presence or absence of 1 nM 5-OP-RU. Sequences of the TCRs are shown in **Fig. S4A**. Cells were subsequently stained with PE-Cy7-conjugated anti-CD69 (BD; 1:100) and PE-labelled anti-MR1 (26.5 clone; BD; 1:100), before analysis on LSRFortessa (BD Biosciences) flow cytometer. Activation of SKW-3-MAIT cells was reflected by the cell-surface CD69 expression. The gating strategy is shown in **Fig. S3B**.

### Human PBMCs

Whole blood was collected in heparin-coated tubes and centrifuged to separate the cellular fraction and plasma using lymphocyte separation solution (d = 1.077) (Nacalai Tesque). PBMCs were stained with anti-human CD3 (HIT3a), anti-human CD161 (HP-3G10), and MR1-Ags-tetramer. For proliferation assay, PBMCs were labeled with CTV (Invitrogen) according to the manufacturer’s instructions. PBMCs were cultured with 5-OP-RU or RF catabolites in RPMI 1640 (Sigma-Aldrich) medium. Seven days after stimulation, PBMCs were analyzed by flow cytometry (Attune NxT flow cytometer, Thermo Fisher Scientific). For primary MAIT activation assays, isolated PBMCs were incubated with RF catabolites for 16 h in the presence or absence of 5-OP-RU. Cells were then stained with anti-human CD69 (BD; 1:200), anti-human CD19 (Biolegend; 1:200), anti-human CD3 (BD; 1:200), anti-human TCR Vα7.2 (Biolegend; 1:200) as well as the viability dye 7AAD for 30 min at RT. Following 2 washes with 2% FACS buffer, cells were incubated with MR1-5-OP-RU tetramers at 1:200 for 30 min before analysis with LSRFortessa (BD Biosciences) flow cytometer. Gating strategy is shown in **Fig. S3C-D**

### Western Blot analysis of intracellular MR1

C1R.MR1 cells were cultured with different RF catabolites for 16 h, at 37□. Samples then treated with or without Endo-H for 1 h incubation at 37□. Cells were lysed and lysate protein concentration was quantified by BCA assay. Equal amounts of samples (40 µg total protein) were separated on SDS-PAGE gels before being transferred onto nitrocellulose membranes and incubated with primary (anti-MR1; Clone:8G3) (McWilliam et al., 2020) then corresponding secondary antibody (IRDye 800CW anti-mouse IgG, Licorbio, Cat# 925-32210). The protein bands were visualized by Licor system.

### Recombinant expression and purification of MR1 and the A-F7 MAIT TCR

*Escherichia coli* BL21(DE3) cells were transformed with plasmids encoding for the extracellular domains of MR1, β2m, and the AF-7 TCRα and TCRβ chains (Kjer-Nielsen et al., 2012; Patel et al., 2013; Wang et al., 2022a). The proteins were then expressed, and the inclusion bodies were purified, cleaned, and used for refolding. For both MR1 and AF-7 TCR, inclusion bodies were refolded through rapid dilution overnight as previously mentioned (Keller et al., 2017). To refold MR1 with ligands, 58 mg MR1 inclusion bodies along with 30 mg of β2m inclusion bodies were placed in a 500 ml of refold buffer consisting of 0.1 M Tris pH 8.0, 5 M of urea, 2 mM EDTA, 2.5 mM oxidized glutathione, 20 mM reduced glutathione and 0.4 M of L-arginine for 16 h at 4° C. Ligands were dissolved in dilute NaOH and added directly into the refolding buffer. For photodegraded RF, 60 mg RF powder was dissolved in 1 L refolding buffer and then exposed to UV light through a UVA lamp for 30 minutes prior to the addition of inclusion bodies. The refolded MR1-ligand complex was dialyzed against 10 mM Tris pH 8.0 overnight and purified using size-exclusion (Superdex 200, GE Healthcare) and anion-exchange (HiTrapQ, GE Healthcare) chromatography techniques (Awad et al., 2020a).

### Thermal stability assay

To investigate the stability of the MR1–RF catabolite complexes, thermal shift assay was performed. Sypro Orange (Sigma) was used as a fluorescent dye to monitor the protein unfolding upon heating. This assay was performed in a real-time detection system (Corbett RotorGene 3000). Each MR1–Ag complex was prepared in 10□mM Tris-HCl (pH 8) and 150□mM NaCl and heated from 28 to 95□°C with a heating rate of 1□°C□min^−1^. The fluorescence intensity was measured (excitation at 530□nm and emission at 610□nm), and the unfolding process was followed in real time. Tm50 represents the temperature at which 50% of the protein was unfolded. All the experiments were performed in triplicates, at three independent times.

### Sample preparation and analysis by LCMS

Refolded MR1 samples containing approximately 10 µg of refolded protein were treated 3:1 with acetonitrile containing 0.1% formic acid. These samples were vortexed and then allowed to stand at RT, protected from light for 10 min, before centrifugation at 15,000 rcf for 3 min to pellet precipitated protein material. The supernatant was transferred to fresh lo-bind Eppendorf tubes and evaporated off using a centrifugal evaporator set to 38°C, reaching dryness after approximately 1 h. The tubes were then reconstituted into an LCMS-matched solvent (mobile phase A; 0.1% formic acid, 2% ACN in Optima water). They were allowed to mix at 37°C for 30 min, prior to another centrifugation step at 15,000 rcf for 3 min, and supernatant transfer to LCMS vials for analysis. Solvent standards for all analytes of interest were also prepared by dilution into the same mobile phase solvent (10 µM) from a 10 mM DMSO stock.

Samples were loaded into an Eksigent NanoLC system which directly injected 5 µL of each sample into a Luna Omega 2 um Polar C18 (100A, LC column 50 x 0.3mm). Gradient chromatography was used with mobile phase B (0.1% formic acid in 80% acetonitrile/Optima water) ramping from 3% to 95% from 2 to 9 min, held for 2 min and returned from 95 to 3% from 11 to 12 min, followed by a 3-min re-equilibration period (15 min total). MS data was acquired using a SCIEX 6600 tripleTOF, scanning MS1 from 100 to 700 m/z, and IDA mode triggering MS2 data acquisition for ions exceeding 100 cps. Dynamic background subtraction and dynamic accumulation functions were used, and up to 10 MS2 scans were acquired per method cycle. Samples were run in both positive and negative modes, with positive mode usually providing meaningful data.

### Crystallization and solving the structures of MR1*01-ligand complexes using MAIT TCR as a crystallization aid

Purified MR1-lumichrome complex was mixed with AF-7 TCR at 1:1 M ratio and incubated for 1 h on ice (Awad et al., 2020a). Because the other RF catabolites couldn’t assist the refolding of MR1 to a reasonable amount in solution, we applied the established MR1 ligand displacement protocol to determine the structure of MR1 loaded with RF, FMF, and lumiflavin (Keller et al., 2017; Salio et al., 2020). Here, purified MR1-empty was concentrated and mixed with AF-7 TCR to a final concentration of 5 mg/ml and the ligand was diluted in this mixture as 1:10 (ligand: MR1) molar ratio. Empty MR1 was prepared as previously described (Keller et al., 2017). The ternary complexes were crystallized in 100 mM Bis-Tris-propane (pH 6.1–6.5), 12 to 18% w/v PEG3350, and 200 mM sodium acetate using hanging drop method. Complex crystals of AF-7 TCR-MR1-lumichrome and MR1-photodegraded-RF complexes were formed within a week. After growth, the crystals were washed in mother liquor supplemented with 12% (v/v) glycerol and flash-frozen in liquid nitrogen. The X-ray diffraction from the ternary crystals was measured at the Australian Synchrotron (McPhillips et al., 2002) and the data are accessed through a Monash local account. The locally processed data were accessed via FileZilla and integrated using XDS package (Kabsch, 2010). We solved the ternary structures at ∼ 1.9-2.2 Å resolution. The AF-7 TCR-MR1-ligand crystal structure was determined by molecular replacement using Phaser (McCoy et al., 2007) and AF-7 TCR-MR1-5-OP-RU structure (PDB code: 6PUC) after removing 5-OP-RU. The structures were initially refined with Phenix.Refine (Liebschner et al., 2019) and then the iterative model was built and improved in Coot^®^ (Emsley and Cowtan, 2004). In all structures, the cut-off for hydrogen bonds, salt bridges and VDW interactions was set at 3.5 Å, 4 Å and 4 Å, respectively. The quality of the structures was confirmed at the Research Collaboratory for Structural Bioinformatics Protein Data Bank Data Validation and Deposition Services website. All presentations of molecular graphics and figures were created with the PyMOL Molecular Graphics System, Version 2.0, Schrödinger, LLC.

### Statistical analysis

An unpaired two-tailed Student’s t test or one-way ANOVA with the Dunnett multiple comparison test was performed for the statistical analyses using GraphPad Prism (Version 9.1.0, GraphPad Software Inc.), using either DMSO or NaOH as a control. Unless otherwise indicated, ‘ns’ refers to ‘not significant’, and Asterisks denote the level of statistical significance (*P<0.05, **P<0.01, ***P<0.005, ****P<0.001). P values were adjusted for multiple comparisons using the Benjamini-Hochberg method (FDR < 0.05) using GraphPad Prism version 9.1.0.

## DATA AND CODE AVAILABILITY

The atomic coordinates of AF-7 TCR-MR1 in complex with riboflavin, FMF, lumiflavin and lumichrome have been deposited in the Protein Data Bank (www.rcsb.org) under accession codes 9O05, 9O06, 9O07 and 9O08, respectively.

## AUTHOR CONTRIBUTIONS

M. Abdelaal: conceptualization, data curation, formal analysis, investigation, methodology, visualization, and writing original draft. J. Deng: investigation, formal analysis and visualization. E. Ito: investigation, formal analysis and visualization. M. McInerney: investigation, formal analysis and visualization. A. W. Purcell: formal analysis and resources. S. Yamasaki: formal analysis and resources. Jose Villadangos: formal analysis and resources. H. E. G. McWilliam: formal analysis, visualization, supervision and resources. N. A. Gherardin: conceptualization, resources and supervision. J. Rossjohn: conceptualization, funding acquisition, project administration, resources, supervision, and writing—original draft and review. W. Awad: conceptualization, formal analysis, funding acquisition, project administration, resources, supervision, and writing—original draft and review

## ACKNOWLEDGEMENTS

This work was supported by the Australian Research Council (ARC; DP250102065), the National Institutes of Health (NIH) RO1 AI148407-01A1. W.A. was supported by Australian ARC Discovery Early Career Researcher Award (DECRA) fellowship (DE220101491) and and Monash FMNHS Future Leader Fellowship. AWP is supported by a NHMRC investigator grant (2016596) and an ARC DP (DP250102065). J.R. is supported by an NHMRC investigator grant (2008981) and an ARC Discovery Project (DP250102065). NAG is supported by an NHMRC EL Investigator Grant (2027058). We would like to thank Jeffrey Mak and David Fairlie for the synthesis of the FMF compound. We would like to thank Dale I. Godfrey and Samuel J. Redmond for their intellectual and technical assistance. We thank the staff at the Monash Macromolecular Crystallization Facility for assistance. This research was undertaken in part using the MX2 beamline at the Australian Synchrotron, part of ANSTO, and used the Australian Cancer Research Foundation (ACRF) detector.

