## Supplemental figures and table for "The antigen presenting molecule MR1 binds riboflavin catabolites"

**This file contains:**

**Fig. S1-S4**

**Table S1**

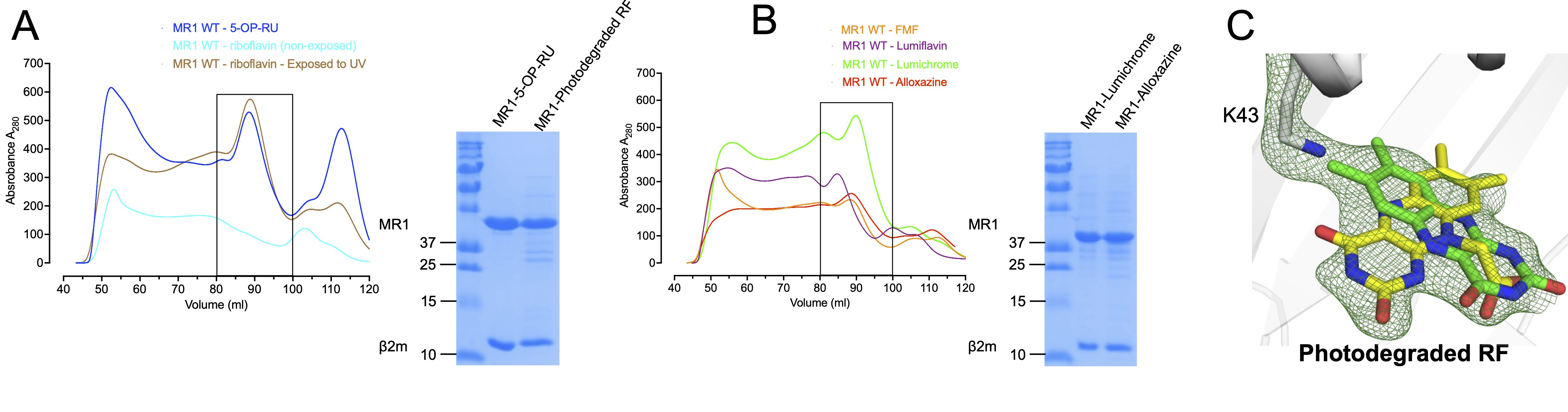

**Figure S1. MR1 refolds with riboflavin catabolites.** Shown is the gel filtration (S200 10/300 GL; GE Healthcare) purification (left) and SDS-PAGE analysis (right) of MR1-β2m binary complexes loaded with **(a)** 5-OP-RU, non-exposed riboflavin, and photodegraded riboflavin, as well as the riboflavin catabolites **(b)** FMF, lumichrome, lumiflavin and Alloxazine. Absorption at 280 nm and volume (ml) are shown on the y- and x-axis, respectively. **(c)** The crystallographic omit maps of UV-treated riboflavin is presented as a F_observed_ – F_calculated_ omit map (green mesh) contoured at 2σ showing the position of the riboflavin degradation product(s) within MR1 cleft connected with Lys-43. The superimposed green and yellow sticks represent lumichrome and carboxymethylflavin, respectively, which have been suggested by mass spectrometry to be the main riboflavin photodegradation catabolites in the refold sample.

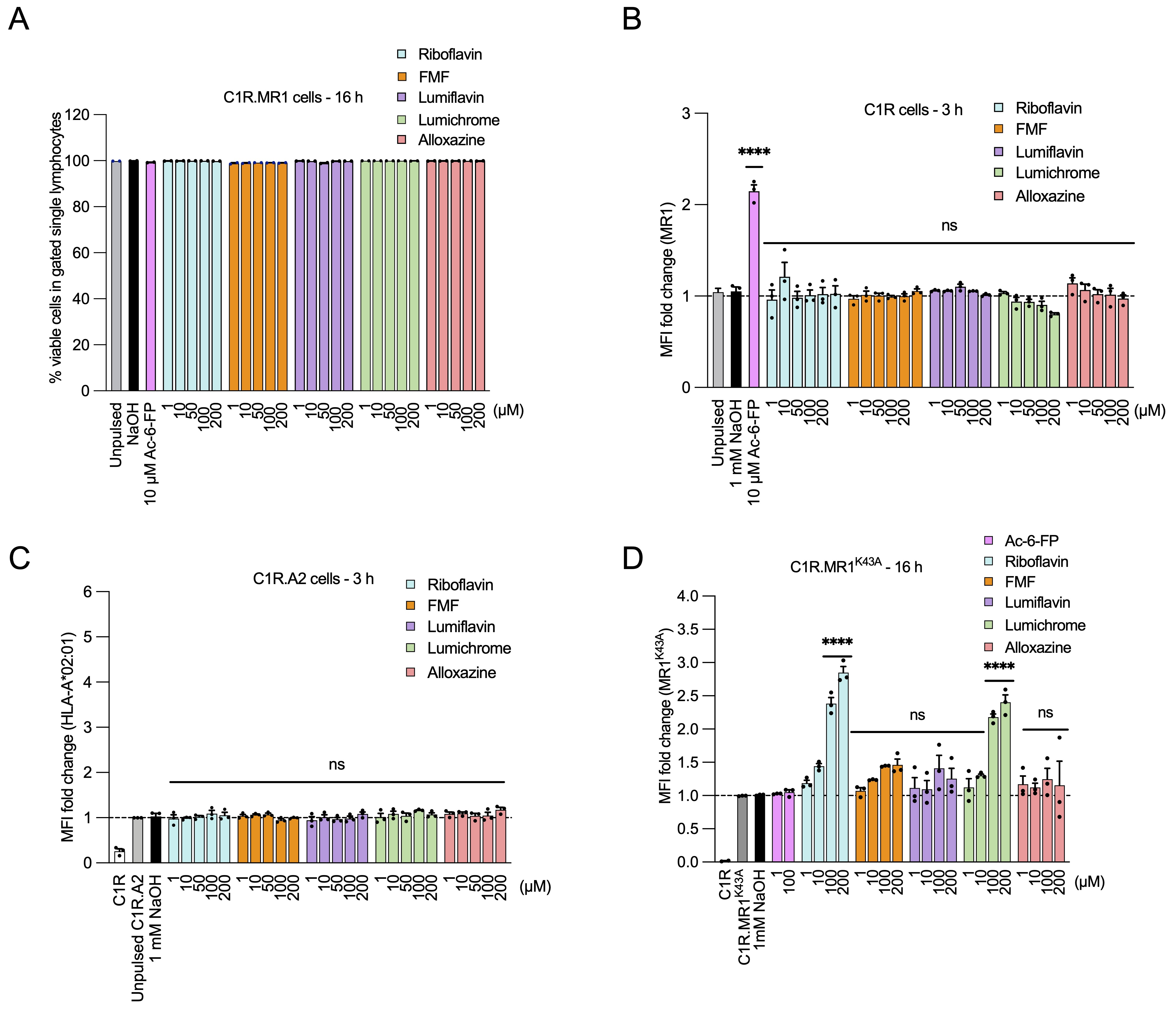

**Figure S2. Responses of different APCs to RF catabolites. (a)** The bar graph shows the frequency of viable cells in single lymphocytes gate for C1R.MR1 cells after 16 h treatment of the indicated ligands. C1R cells expressing **(b)** wild-type level of MR1*01, or overexpressing **(c)** HLA-A*02:01 or **(d)** MR1.K43A mutant were incubated for the indicated periods with titrated quantities of ligand followed by flow cytometry. The C1R and C1R.A2 cells were stained with biotinylated 8F2F9 and W6/32, respectively; followed by PE-labelled streptavidin. In all experiments, the MR1 and HLA-A*02:01 surface expression are depicted as a fold change from basal surface expression upon incubation with the vehicle (1 mM NaOH). Each column represents the average of at least three independent experiments performed in duplicate with standard error (SEM) represented by error bars. One-way ANOVA statistical analysis was performed for all samples followed by Dunnett multiple comparison test using NaOH as a control (ns; not significant, * *p* <0.05, ***p* < 0.01, ****p* < 0.001, *****p* < 0.0001).

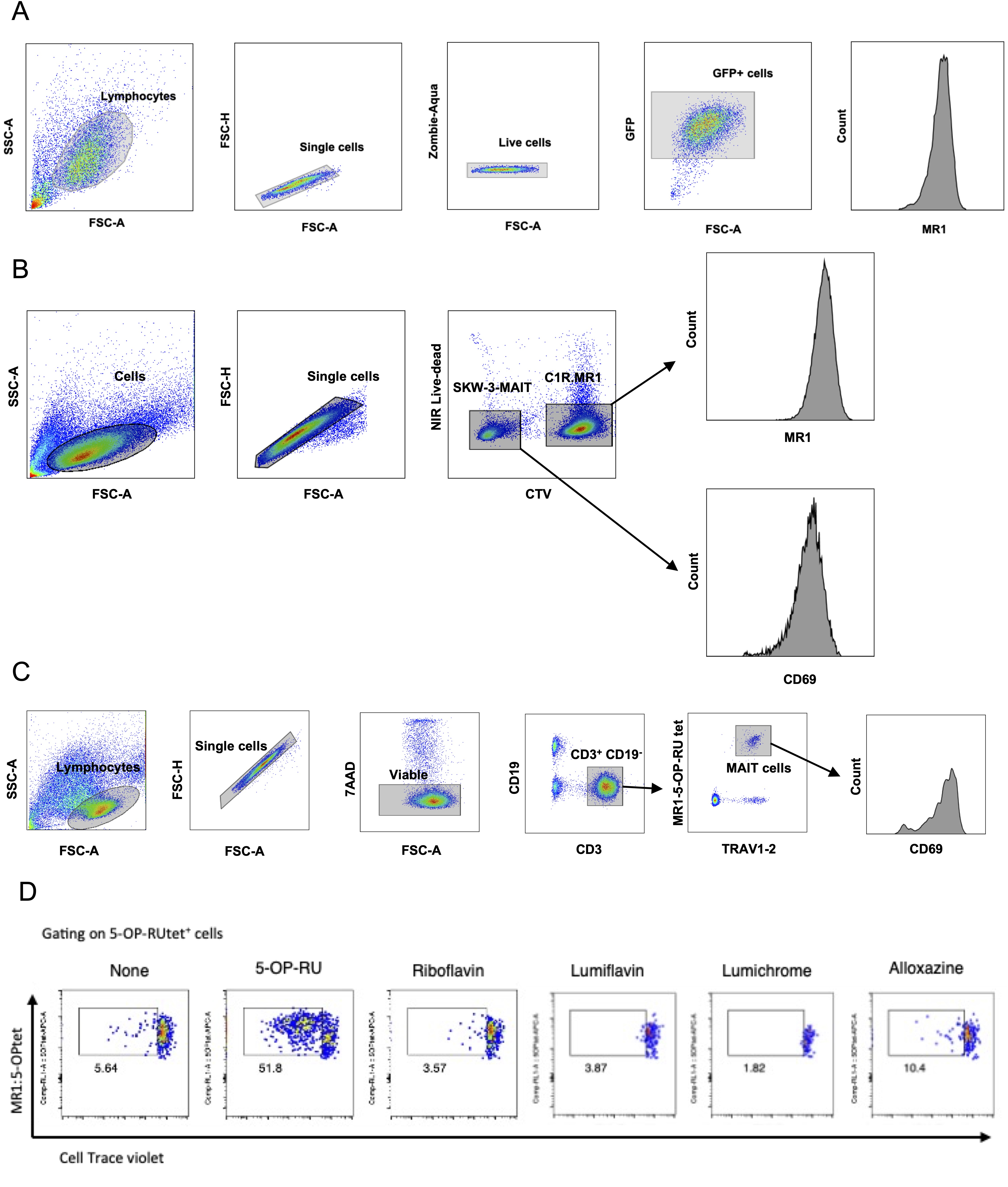

**Figure S3. Gating strategies of flow cytometry experiments**. Shown are the gating strategies used in **(a)** MR1 upregulation experiments, **(b)** MAIT activation/inhibition experiments and **(c)** PBMCs experiments. (**d**) Representative flow cytometry dot plots of proliferating CD3^+^MR1−5-OP-RU tet^+^CTV^lo^ MAIT cells related to **Fig. 5A**.

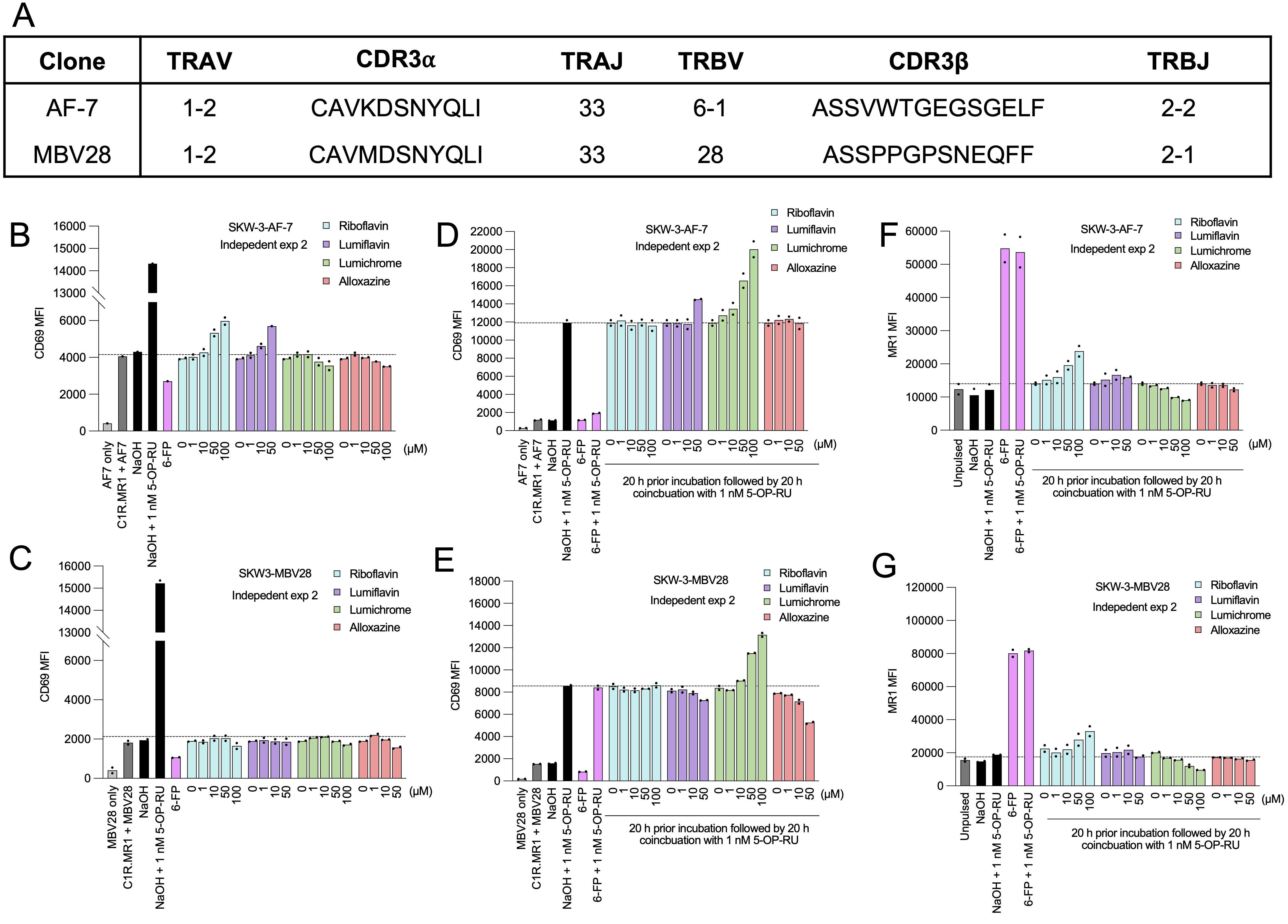

**Figure S4. Biological replicates and TCR sequences of the MAIT activation experiments. (a)** Shown are the sequences of MAIT TCRs used in the MAIT activation experiments as described in the Materials and Methods section. The bar graphs show a second biological replicate for the CD69 expression data shown in **Fig. 4** for two separate SKW-3-AF-7 and SKW-3-MBV28 cell lines in the absence **(b-c)** or presence of 5-OP-RU **(d-e)** after preincubation with titrated quantities of indicated ligands followed by harvesting, staining and flow cytometric analysis. The MR1 expression on C1R.MR1 cell surface matching the (d-e) 5-OP-RU co-incubation experiments is shown in **(f-g)**.

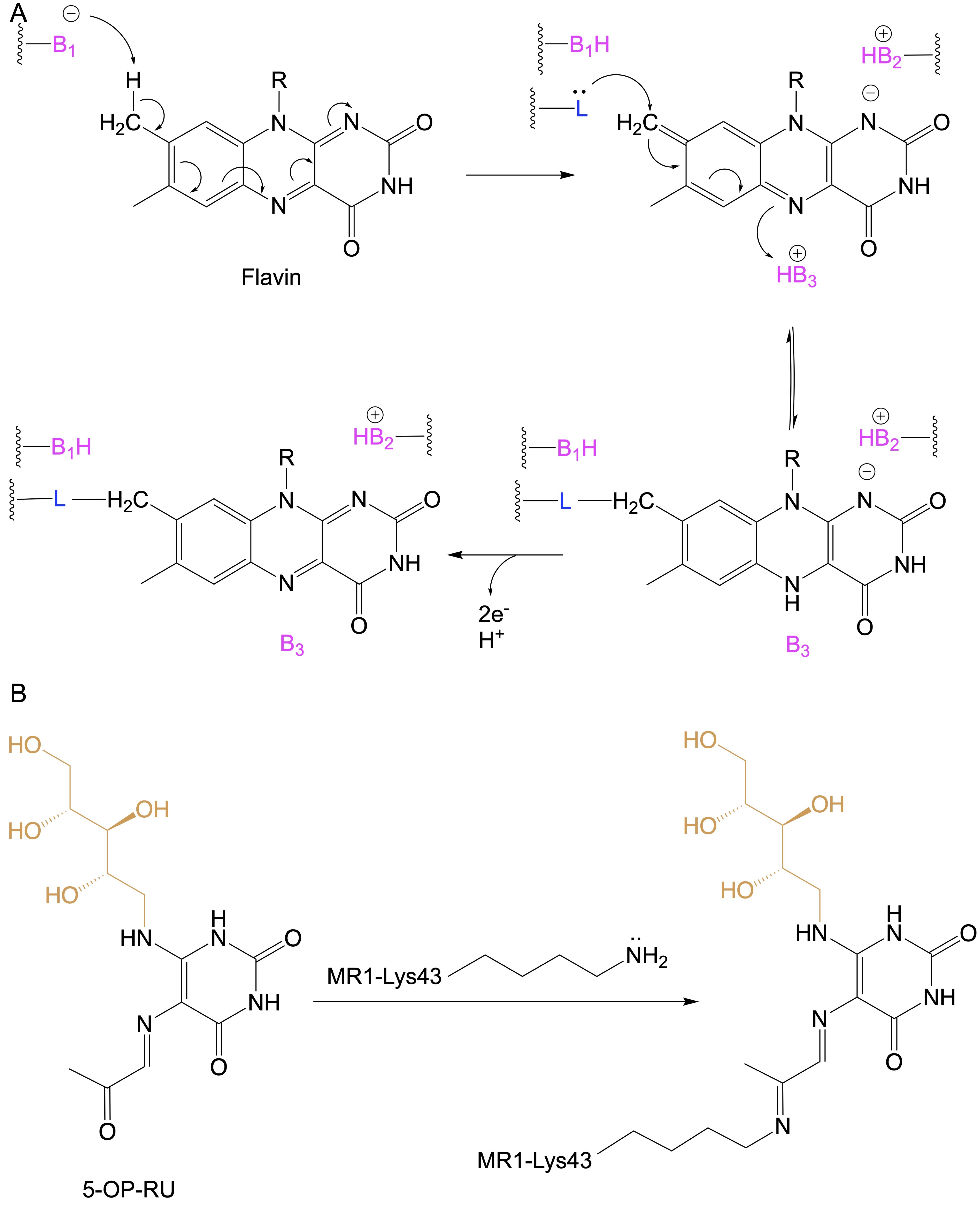

**Figure S5. Comparison between the flavin bond and Schiff base formation.** **(a)** Shown is the general proposed mechanism for the formation of covalent flavin-protein bond at the isoalloxazine ring, adapted from (Heuts et al., 2009). Here, B_1_, B_2_ and B_3_ are basic side chains while L^-^ is the nucleophilic side chains. **(B)** The Schiff base is formed through imine condensation between the carbonyl group of a ligand (here, 5-OP-RU) and the amino group of MR1-Lys43.

**Table S1. Data collection and refinement statistics.**

|  | **A-F7 TCR-MR1-Lumichrome** | **A-F7 TCR-MR1-Riboflavin** | **A-F7 TCR-MR1-Lumiflavin** | **A-F7 TCR-MR1-FMF** |
| --- | --- | --- | --- | --- |
| **Resolution range** | 45.43 - 2.2 (2.22 - 2.20) | 47.01 - 1.95 (1.97 - 1.95) | 49.32 - 1.97 (1.99 - 1.97) | 48.16 - 2.0 (2.02 - 2.00) |
| **Space group** | C 1 2 1 | C 1 2 1 | C 1 2 1 | C 1 2 1 |
| **Unit cell**  **a, b, c (Å)**  **α, β, γ (°)** | 216.43 69.73 143.13  90 104.38 90 | 217.38 70.77 143.63  90 104.70 90 | 212.71 69.29 141.17 90 103.48 90 | 217.42 70.32 143.28  90 104.63 90 |
| **Total reflections** | 204792 (6827) | 295931 (9789) | 278045 (9097) | 279863 (9326) |
| **Unique reflections** | 104865 (3501) | 152073 (5066) | 141041 (4650) | 141294 (4721) |
| **Multiplicity** | 2.0 (2.0) | 1.9 (1.9) | 2.0 (2.0) | 2.0 (2.0) |
| **Completeness (%)** | 99.45 (99.37) | 98.67 (98.01) | 99.66 (99.51) | 99.52 (99.43) |
| **Mean I/sigma(I)** | 8.60 (1.58) | 14.68 (1.69) | 11.03 (1.70) | 10.29 (1.24) |
| **Wilson B-factor (Å^2^)** | 41.26 | 31.88 | 37.75 | 39.30 |
| **R-merge** | 0.04655 (0.4859) | 0.02797 (0.47) | 0.02892 (0.4591) | 0.02778 (0.4551) |
| **R-meas** | 0.06584 (0.6872) | 0.03956 (0.6647) | 0.04089 (0.6493) | 0.03929 (0.6437) |
| **R-pim** | 0.04655 (0.4859) | 0.02797 (0.47) | 0.02892 (0.4591) | 0.02778 (0.4551) |
| **CC1/2** | 0.997 (0.802) | 0.999 (0.743) | 0.999 (0.618) | 0.999 (0.838) |
| **CC*** | 0.999 (0.943) | 1 (0.923) | 1 (0.874) | 1 (0.955) |
| **R-work** | 0.1663 (0.2742) | 0.1660 (0.2737) | 0.1574 (0.2417) | 0.1739 (0.3418) |
| **R-free** | 0.2190 (0.3404) | 0.1657 (0.2979) | 0.1920 (0.2682) | 0.2114 (0.3838) |
| **Number of non-hydrogen atoms** | 14146 | 14765 | 14486 | 14166 |
| **macromolecules** | 13053 | 13043 | 13031 | 12969 |
| **ligands** | 67 | 100 | 45 | 110 |
| **solvent** | 1026 | 1622 | 1410 | 1087 |
| **Protein residues** | 1607 | 1606 | 1606 | 1606 |
| **RMS (bonds) (Å)** | 0.007 | 0.006 | 0.007 | 0.007 |
| **RMS (angles)** | 0.86 | 0.82 | 0.85 | 0.81 |
| **Ramachandran favored (%)** | 97.85 | 98.23 | 98.29 | 97.85 |
| **Ramachandran allowed (%)** | 2.09 | 1.71 | 1.71 | 2.15 |
| **Ramachandran outliers (%)** | 0.06 | 0.06 | 0.00 | 0.00 |
| **Rotamer outliers (%)** | 1.49 | 2.12 | 1.84 | 2.07 |
| **Average B-factor** | 48.17 | 42.86 | 42.98 | 53.82 |
| **macromolecules** | 48.00 | 42.31 | 42.48 | 53.83 |
| **ligands** | 46.29 | 37.96 | 32.09 | 45.23 |
| **solvent** | 50.43 | 47.63 | 47.97 | 54.52 |

*Statistics for the highest-resolution shell are shown in parentheses.
